# Simultaneous human intracerebral stimulation and HD-EEG: ground-truth for source localization methods

**DOI:** 10.1101/2020.02.14.948984

**Authors:** Ezequiel Mikulan, Simone Russo, Sara Parmigiani, Simone Sarasso, Flavia Maria Zauli, Annalisa Rubino, Pietro Avanzini, Anna Cattani, Alberto Sorrentino, Steve Gibbs, Francesco Cardinale, Ivana Sartori, Lino Nobili, Marcello Massimini, Andrea Pigorini

## Abstract

Precisely localizing the sources of brain activity as recorded by EEG is a fundamental procedure and a major challenge for both research and clinical practice. Even though many methods and algorithms have been proposed, their relative advantages and limitations are still not well established. Moreover, these methods involve tuning multiple parameters, for which no principled way of selection exists yet. These uncertainties are emphasized due to the lack of ground-truth for their validation and testing. Here we provide the first open dataset that comprises EEG recorded electrical activity originating from precisely known locations inside the brain of living humans. High-density EEG was recorded as single-pulse biphasic currents were delivered at intensities ranging from 0.1 to 5 mA through stereotactically implanted electrodes in diverse brain regions during pre-surgical evaluation of patients with drug-resistant epilepsy. The uses of this dataset range from the estimation of in vivo tissue conductivity to the development, validation and testing of forward and inverse solution methods.

## Background & Summary

Electroencephalography (EEG) records brain electric potentials through electrodes placed on the scalp. This technique has a relatively low spatial resolution as compared to others (i.e. intracranial EEG, functional Magnetic Resonance Imaging, etc.), mainly due to volume-conduction induced spatial averaging^1,2^. However, in the last decades, a plethora of methods have been developed aimed at reconstructing the sources of the activity recorded from the scalp^3^. The procedure involves, first, creating a model of how electrical currents propagate from their origin to the recording electrodes, the so-called forward problem; and second, creating a model of the plausible locations and intensities of the current sources that gave rise to the recorded activity, the so-called inverse problem. Many methods exist for solving each of these two problems. Forward models range from a single spherical shell to a detailed reconstruction of the various tissues and geometrical characteristics of specific individuals (for a review see^4^). Likewise, inverse models range from estimating a single dipole at a fixed pre-established location to calculating thousands of them distributed following the cortical geometry of a particular subject (for a review see^5^).

Despite being widely used, validating and comparing these methods remains a controversial issue due to the lack of ground-truth data. Most methods’ validations rely on simulations in order to assess their accuracy and robustness^6,7^. That is, simulated electrical activity is placed inside a realistic volume-conductor model and projected onto the scalp surface in order to be used as input data for source localization algorithms, which are then tested on their ability to reconstruct the origins of these signals. Another common methodology is to try localizing functional activity whose origins are inferred from other imaging modalities^8^ (i.e. fMRI during somatosensory stimulation). However, simulations lack realism and cross-modal functional mapping lacks spatial precision and can introduce relative biases in spatial arrangement due to the different nature of the signals.

A fundamental element to fill this gap could be offered by stereo-electroencephalography (sEEG), obtained from drug-resistant epileptic patients using stereotactically implanted electrodes. Once surgically implanted, patients are monitored continuously for several days to have one or more seizures recorded. During this time, sessions of intracortical stimulation are performed in order to induce habitual seizures and to provide a map of the physiological functions of the implanted sites^9–14^. This procedure implies that a brief current pulse is injected between two adjacent leads, producing an electrical artifact whose localization can be accurately determined. When combined with simultaneous scalp EEG, this procedure is capable of generating real data of scalp recorded electrical signals originating from precisely known locations inside the human brain, and thus represents an ideal benchmarking scenario for validating and comparing both forward and inverse solution methods.

In line with this, the aim of this paper is to provide a consistent dataset of high-density scalp EEG recordings performed during the stimulation of intracortical leads. It contains the anonymized MRIs necessary to build forward models, the surfaces and forward models created using the subjects’ original MRIs, the spatial and anatomical information of the stimulated sites, and EEG data from 256 channels with digitized positions. As a further element, stimulations were performed at different current intensities, so as to favor not only a comparative performance across different topographical regions, but also an estimation of the role that the intensity of a source activity plays in its localization accuracy. The value of this dataset is also increased by the dense sampling of the scalp, which allows spatial down-sampling procedures to test the performance of inverse solution algorithms under a montage-dependent perspective.

In order to demonstrate the validity and wide range of possible uses of this dataset, we performed three different analysis. First, we tested the performance of three widely used inverse solution methods, employing various montages and parameters’ configurations, and tested the best reachable performance. Second, we examined how misselection of parameters affected localization accuracy. Finally, we evaluated how different MRI anonymization procedures influence source localization results.

To the best of our knowledge, this would be the first dataset providing the neuroscientific and technical community with ground truth to validate the efficacy of forward and inverse solutions on EEG data, and to systematically evaluate the factors mostly contributing to the overall process accuracy.

## Methods

### Participants

Seven subjects (F = 4) participated in the study (<u>*X*</u>age = 35.1; *sd* age= 5.4). A total of 61 sessions were obtained (<u>*X*</u>sessions per subject = 8.71; *sd* sessions per subject = 2.65). All subjects were patients undergoing intracranial monitoring for pre-surgical evaluation of drug-resistant epilepsy. All of them provided their Informed Consent before participating, the study was approved by the local Ethical Committee (protocol number: 463-092018, Niguarda Hospital, Milan, Italy) and it was carried out in accordance with the Declaration of Helsinki.

**Table 1.** Participants’ demographic and clinical information. Subject code, sex, age at the time of evaluation, language dominant hemisphere, epileptogenic zone and pharmacology (morning/noon/night intakes; when only one value is present it corresponds to a single day intake).

| subject | sex | age | hemispheric dominance | epileptogenic zone | pharmacology |
| --- | --- | --- | --- | --- | --- |
| sub-01 | M | 37 | R | Right fronto-central | Carbamazepine: 400/0/400 mg;<br>Lacosamide: 150/0/150 mg |
| sub-02 | F | 39 | R | Left temporo-mesial | Carbamazepine: 400/200/400 mg;<br>Levetiracetam: 1000/750/1000 mg;<br>Clobazam: 0/0/10 mg |
| sub-03 | M | 35 | R | Left temporo-mesial | Levetiracetam: 1500/0/1500 mg;<br>Lacosamide: 200/0/200 mg;<br>Carbamazepine: 800/0/600 mg |
| sub-04 | M | 44 | R | Right temporo-mesial | Carbamazepine: 400/200/400 mg;<br>Perampanel: 6/0/0 mg |
| sub-05 | F | 28 | R | Left temporo-mesial | Carbamazepine: 600/0/600 mg;<br>Perampanel: 6 mg |
| sub-06 | F | 32 | R | Right temporo-mesial | Carbamazepine: 400/0/400 mg;<br>Zonisamide: 100/0/100 mg; Clobazam:<br>0/0/10 mg |
| sub-07 | F | 31 | R | Superior temporal gyrus | Carbamazepine: 600/600 mg;<br>Topiramate: 50/100 mg |

### Electrical stimulation

Intracranial shafts were implanted using a robotic assistant (Neuromate; Renishaw Mayfield SA), with a workflow detailed elsewhere^13^. The position of the electrodes was decided exclusively following clinical needs. Electrical currents were delivered through platinum-iridium semiflexible multi-contact intracerebral electrodes (diameter: 0.8 mm; contact length: 2 mm, inter-contact distance: 1.5 mm; Dixi Medical, Besançon, France). Single-pulse biphasic currents lasting 0.5 ms were delivered at intensities ranging from 0.1 to 5 mA (number of sessions: 0.1 mA = 22; 0.3 mA = 17, 0.5 mA = 8; 1 mA = 9; 5 mA = 5) through pairs of adjacent contacts by a Nihon-Kohden Neurofax-100 system. The stimulation frequency (i.e. number of pulses per second) was of 0.5 Hz when stimulating at 1 and 5 mA and 1 Hz otherwise (with the exception of 3 sessions at 1 mA on which the stimulation frequency was 1 Hz). A total of 60 trials were obtained from each stimulation site when stimulating at 0.1, 0.3 and 0.5 mA, and a total of 40 when stimulating at 1 and 5 mA (Figure 2).

**Figure 1.**
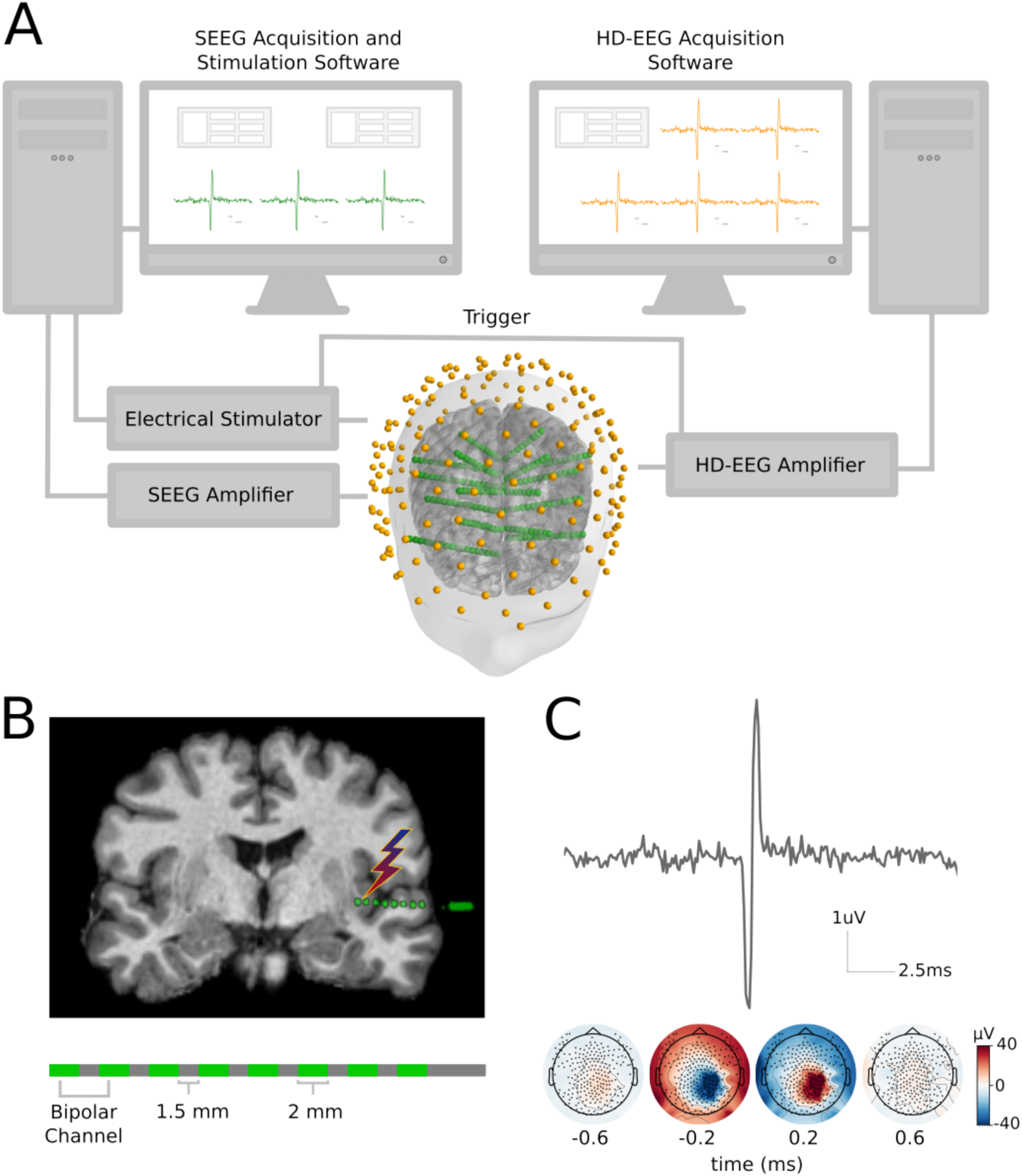
Illustration of the experimental setup. A) Depiction of the stimulation and acquisition systems’ temporal synchronization and spatial co-registration. B) *Top:* example of an intracerebral shaft containing eight contacts coregistered with the subject’s MRI. *Bottom:* Illustration of an intracranial shaft. C) *Top:* Example of a stimulation artifact recorded by a scalp EEG channel. *Bottom:* Scalp EEG topographies at the time of the stimulation onset.

**Figure 2.**
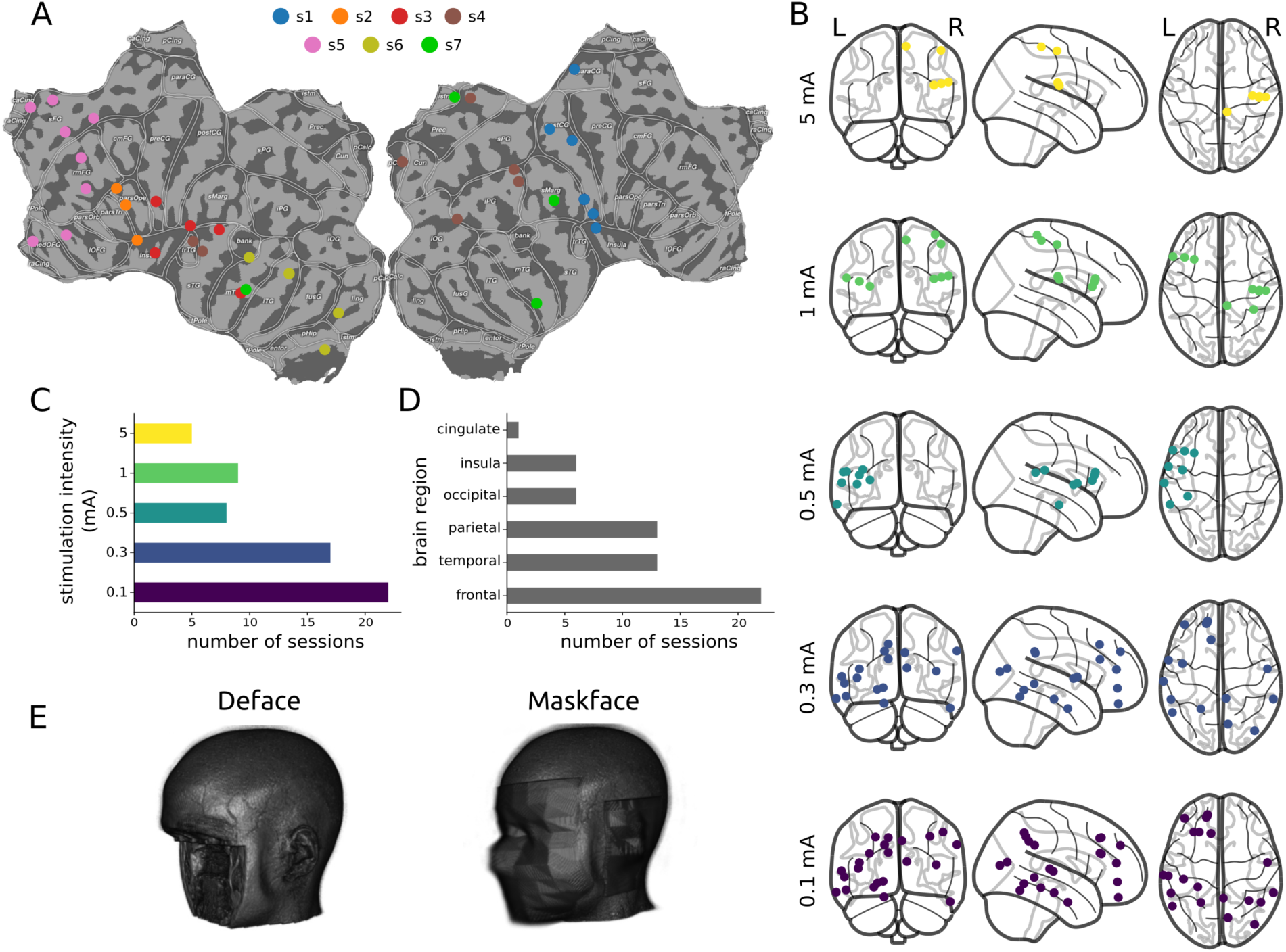
Dataset description. A) Flatmap of stimulation sites by subject. B) Location of stimulation sites by stimulation intensity. C) Number of sessions by stimulation intensity. D) Numbersessions by brain lobe. E) Example of the anonymization methods. The MRI shown belongs to an open dataset^34^ as it was not possible to show images of the participants of the study due to privacy issues.

### EEG Recordings

EEG signals were recorded from 256 channels (Geodesic Sensor Net; HydroCel CleanLeads) sampled at 8000 Hz with an EGI NA-400 amplifier (Electrical Geodesics, Inc; Oregon, USA), using a custom-built acquisition software written in C++ and Matlab, based on EGI’s AmpServerPro SDK. All software filters were disabled during acquisition. The spatial locations of EEG electrodes and anatomical fiducials were digitized with a SofTaxicOptic system (EMS s.r.l., Bologna, Italy), coregistered with a pre-implant MRI (Achieva 1.5 T, Philips Healthcare).

### Electrode localization

The location of the intracranial electrodes was assessed registering the post-implant CT (O-arm 1000 system, Medtronic) to the pre-implant MRI by means of the FLIRT software tool^15^. The position of every single lead was assessed with respect to the MRI using Freesurfer^16^, 3D Slicer^17^ and SEEG assistant^18^. When the pre-implant MRI and the EEG digitization MRI were not the same, contacts positions were transformed from the SEEG space to the EEG space using an affine transformation between MRIs calculated employing the ANTs software^19^. Normalized contacts’ coordinates were estimated by performing a non-linear registration between the subject’s skull stripped MRI and the skull-stripped MNI152 template^20^ (ICBM 2009a Nonlinear Symmetric) using ANTs’ *SyN* algorithm. Contact positions were plotted on a flatmap of the MNI152 template built using Pycortex^21^, by projecting each contact’s coordinates to the closest vertex of the brain surface reconstruction. The accuracy of the normalization procedure was verified by visual inspection.

### Data preprocessing

Raw data were imported and preprocessed in Python employing custom-built scripts and the MNE software^22,23^. Continuous data were high-pass filtered at 0.1 Hz (FIR filter; zero phase; Hamming window; automatic selection of length and bandwidth). Data from two subjects (sub-05 and sub-07) were also notch filtered at 50, 100, 150 and 200 Hz (FIR filter; zero phase; Hamming window; bandwidth = 0.1 and automatic length selection) due to considerable line noise. Bad channels were identified by visual inspection (i.e. flat channels, presence of artifacts, etc.). Next, epochs were generated from -300 ms to 50 ms with respect to the stimulation electrical artifact and baseline corrected (mean subtraction method, from -300 ms to -50 ms). The baseline period was specifically chosen to avoid any possible contamination by cortico-cortical evoked responses from previous trials, even with the fastest stimulation frequency^24^. Bad epochs were identified by visual inspection and rejected. Given that EGI’s trigger channel is sampled at 1000 Hz, which introduced jitter between the onset of the trigger and the onset of the stimulation, epochs were fine-aligned by matching the peaks of the stimulation artifacts within sessions. All good epochs were saved in MNE’s *fif* format in the interval between -250 and 10 ms and subsequently converted to BIDS format^25,26^using custom code based on the MNE-BIDS^27^package.

### Source localization

The source localization procedure was carried out using the MNE software. Surface reconstructions were obtained with Freesurfer and a 3-layer Boundary Element Method (BEM) model was created with 5120 triangles and conductivities set to 0.3, 0.006 and 0.3 S/m, for the brain, skull and scalp compartments respectively. Source spaces were created with 4098 sources per hemisphere. Epochs were re-referenced to the average of all good channels and covariance was estimated with automated method selection^28^.Subsequently, epochs were averaged and cropped from -2 to 2 ms with respect to the stimulation artifact. Inverse solutions were calculated with three different methods: Minimum Norm Estimate (MNE), dynamic Statistical Parametric Maps (dSPM) and exact Low Resolution Electromagnetic Tomography (eLORETA) ^5,29–31^.

Various parameter configurations were assessed. The regularization parameter was set as 1 / SNR^2^ with SNR set to 1, 2, 3, and 4. The depth and loose weighting parameters varied between 0.1 and 1 in 0.1 steps. Four different EEG montages were tested: all good channels, and channels corresponding to EGI’s 128, 64 and 32 montages. When a channel selected for the subsampled montage was marked as bad, we replaced it by its closest neighbour. A total of 4800 solutions were calculated for each session.

The Euclidean distance between the coordinates of the center of the pair of stimulating contacts and the coordinates of the maximal activation in the source estimates were computed as well as the distance on each spatial axis (left-right, anterior-posterior and inferior-superior) as measures of accuracy. We then computed the best solution across all montages and parameter’s configurations. Finally, we computed the proportion of sessions on which each method and montage subsampling reached the distance of the best solution.

### MRI anonymization

MRIs were anonymized employing two different tools: Pydeface (https://github.com/poldracklab/pydeface) and MaskFace^32^(Figure 2.E). In order to investigate the influence on source localization results of the geometrical distortions induced by the anonymization procedures, we recreated the forward-models with the anonymized MRIs and computed the inverse solutions of all the parameters’ configurations that reached the minimum distance of each session. We then compared the distances to the stimulation sites obtained with the anonymized MRIs with the ones obtained with the original ones.

### Data Records

The data is available at the Human Brain Project platform (https://kg.humanbrainproject.eu/instances/Dataset/f557d71e-fe11-43d7-8225-7c2d432f34b9; DOI: 10.25493/NXN2-05W). The dataset comprises high density-EEG data from a total of 61 sessions, obtained from 7 subjects. In addition, it includes the spatial locations of the stimulating contacts in native MRI-space^33^, MNI152-space and Freesurfer’s surface-space, and the digitized positions of the 256 scalp EEG electrodes. It also contains the BEM, pial and inflated surface reconstructions created with the subjects’ original MRIs, as well as the source-spaces and forward-models from them derived. Furthermore, it includes the anonymized MRIs of each subject.

## Technical Validation

### Methods, montages and parameters

The minimum distance between the stimulation sites and the location of the maximum current values was between ∼2 and ∼20 mm when optimal parameters were selected (<u>*X*</u>minimum distance = 6.71 mm; sd minimum distance = 4.15, *min* minimum distance = 2.32, *max* minimum distance = 19.85; Figure 3.A). Instead, when all parameters’ configurations were considered, the distance between the stimulation site and the location of the maximum current values was generally between ∼ 2 mm and ∼ 50 mm (Figure 3.B & Figure 3.C). The proportion of sessions on which each method reached the minimum distance was of 0.14 for MNE, 0.42 for dSPM and 0.57 for eLORETA (Figure 3.D). The proportion of sessions on which each montage reached the minimum distance was 0.26 for all good channels, 0.39 for 128 channels, 0.37 for 64 channels and 0.39 for 32 channels (Figure 3.E). The differences between the stimulation site and the location of the maximum current value of the solutions that reached the best solution for each session were approximately centered around zero and symmetrical across the three spatial axes (L-R, A-P, I-S).

**Figure 3.**
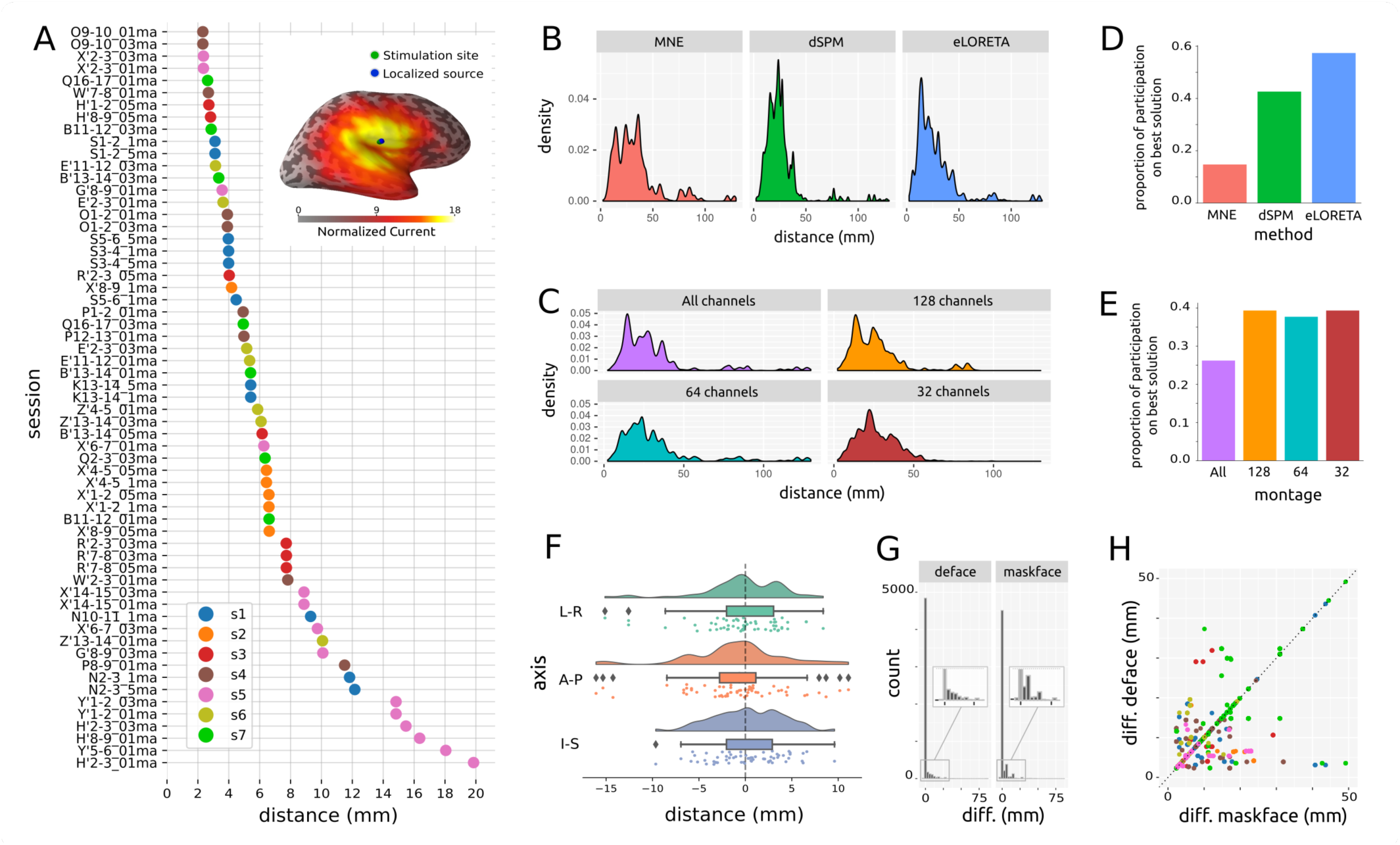
Validation. A) Distance between stimulation site and location of the maximum current value of the best solution for each session. Colors represent subjects. *Insert*: Position of the stimulated site, localized source and estimated current values for a representative session. B) Density plot of distances between the stimulation site and the location of the maximum current value across all parameters’ combinations by inverse solution method. C) Density plot of distances between the stimulation site and the location of the maximum current value across all parameters’ combinations by montage sub-sampling. D) Proportion of sessions on which each inverse solution method reached the minimum distance. E) Proportion of sessions on which each montage subsampling reached the minimum distance. F) Density plot, boxplot and scatterplot of the difference between stimulation site and location of maximum activation of best solution on each session by spatial axis (L-R: left-right; A-P: anterior-posterior; I-S: inferior-superior). G) Histogram of differences between the distance of the stimulation site and the location of the maximum current value between the inverse solutions computed with the original MRI and those computed with the anonymized MRIs. *Insert:* zoom-in on the marked section of the histogram. H) Scatterplot of distance between stimulation site and maximum current value between anonymization methods for each parameters’ configurations that reached the minimum distance for each session. Colors represent subjects as color-coded in panel A.

### MRI anonymization

The distance between the stimulation sites and the location of the maximum current values remained equal in a relatively large number of solutions when employing the anonymized MRIs for the calculation of the forward models (Figure 3.G), with both anonymization methods (% equal deface = 0.88; % equal maskface = 0.82). However, a number of them proved to produce different results. Moreover, the solutions on which the results were different from those obtained with the original MRIs were not the same across anonymization methods (Figure 3.H).

## Usage Notes

The data are provided in BIDS format and contains all the necessary information to allow researchers to perform their analysis on any software. However, please note that, at the time of publication of this article, the BIDS specification for Common Electrophysiological Derivatives has not been established yet and therefore the dataset structure might not be compatible out-of-the-box with all software. However, adjusting the structure for specific purposes should be straight-forward and, importantly, once the specification will be published, we will update the database in order to conform to it. Interactive scripts of usage demonstration are provided as part of the repository accompanying this article.

This dataset has multiple potential uses, for instance: estimating in-vivo tissue conductivities; evaluating the impact of different forward-models on inverse solutions; developing, validating and testing different inverse solution methods; studying interactions between forward and inverse solution methods; performing linear combinations of stimulation sessions in order to test the ability of diverse methods to retrieve the correct sources; etc.

It is worth mentioning that the artifacts generated by intracranial stimulation are non-physiological, therefore generalization of results to physiological signals should be done conscientiously. Also, in some cases, the tails of the intracranial shafts, which protruded from the scalp, precluded the contact with the skin of a number of EEG electrodes. Nevertheless, the analysis performed revealed good localization accuracy, demonstrating that this was not an issue. Another limitation corresponds to the fact that anatomical areas sampled tend to be clustered within subjects, which should be taken into consideration when performing topographical analysis. However, the dataset will be extended with data from new subjects in the future, which will provide a more comprehensive spatial coverage and allow more detailed spatial analyses.

## Code availability

Usage demonstration scripts and the code used for the preparation, pre-processing and technical validation of the dataset are publicly available at https://github.com/iTCf/mikulan_et_al_2019.

## Acknowledgments

This research has received funding from the European Union’s Horizon 2020 Framework Programme for Research and Innovation under the Specific Grant Agreements No. 720270 and No. 785907 (Human Brain Project SGA1 and SGA2; to M.M. and P.A.); and from the Italian Ministry of Health, Targeted Research Grant No. RF-2010-2319316 (to L.N.).

## Author Contributions

E.M., S.S., P.A., L.N., M.M. and A.P. conceived and designed the study. E.M., S.R., S.P., F.Z., A.R., I.S. and A.P. carried out the experiments. F.C., I.S., & L.N. took care of the patients’ clinical management. E.M. analyzed the data, created the figures and curated the database. E.M., S.R., S.P., S.S., F.M.Z., P.A., A.C., A.S., S.G., F.C., I.S., L.N., M.M. and A.P. wrote the article.

